# Cortical myosin minifilaments orchestrate the arrangement of microridge protrusions on epithelial cell surfaces

**DOI:** 10.1101/2020.10.22.351312

**Authors:** Aaron P. van Loon, Ivan S. Erofeev, Andrew B. Goryachev, Alvaro Sagasti

**Affiliations:** Department of Molecular, Cell and Developmental Biology, and Molecular Biology Institute, University of California, Los Angeles, Los Angeles, CA; Centre for Synthetic and Systems Biology, School of Biological Sciences, University of Edinburgh, Edinburgh, UK

## Abstract

Actin-based protrusions vary in morphology, stability, and arrangement on cell surfaces. Microridges are laterally-elongated protrusions arranged in maze-like patterns on mucosal epithelial cells that rearrange dynamically by fission and fusion. To characterize how microridges mature and investigate the mechanisms driving fission and fusion, we imaged microridges in the maturing skin of zebrafish larvae. After their initial development, microridges continued to lengthen and microridge alignment became increasingly well ordered. Imaging F-actin and Non-Muscle Myosin II (NMII) revealed that microridge fission and fusion were associated with local NMII activity in the apical cortex. Inhibiting NMII blocked rearrangements, reduced microridge density, and altered microridge spacing. High-resolution imaging revealed that individual cortical NMII minifilaments are tethered to protrusions, often connecting adjacent microridges. NMII minifilaments connecting the ends of microridges fused them together, whereas minifilaments oriented perpendicular to microridges severed them or pulled them closer together. Our findings demonstrate that as cells mature, microridges continue to remodel and form an increasingly orderly arrangement through a process orchestrated by cortical NMII contraction.

## INTRODUCTION

Cells create diverse actin-based protrusions to carry out a wide variety of functions. Not only do protrusions vary in shape and size, but also in persistence, dynamics, and their relative arrangement on cells. For example, lamellipodia extend and retract within seconds or minutes (Giannone et al., 2007), whereas invadopodia persist for hours (Murphy and Courtneidge, 2011), and stereocilia are stable throughout an animal’s life (Narayanan et al., 2015; Zhang et al., 2012). The stability and plasticity of protrusions depends on the regulation of their constituent actin filaments, but those regulatory mechanisms vary. For example, despite the fact that microvilli maintain a relatively stable height, actin filaments within them are constantly turned over (Loomis et al., 2003; Meenderink et al., 2019; Tyska and Mooseker, 2002). By contrast, the stability of stereocilia reflects extreme stability of their actin filaments, which persist for months (Narayanan et al., 2015; Zhang et al., 2012). The motility and relative arrangement of protrusions on cells are also regulated by diverse mechanisms. For instance, microvilli move rapidly and independently on cell surfaces (Meenderink et al., 2019), but form stable clusters by establishing protocadherin-based connections at their tips (Crawley et al., 2014; Meenderink et al., 2019). Stereocilia, on the other hand, form highly stable and stereotyped arrangements on cells, and their orientation is strictly dictated by planar cell polarity (Tarchini and Lu, 2019). Identifying mechanisms regulating the stability and arrangement of protrusions is critical to understanding how cell surfaces acquire diverse morphologies and adapt to tissue-level changes.

Microridges are laterally-elongated protrusions arranged in elaborate patterns on the apical surfaces of mucosal epithelial cells (Depasquale, 2018). Neighboring microridges tend to align parallel to one another, filling cell surfaces in maze-like arrangements. Although microridges are less studied than other protrusions, recent work in zebrafish periderm cells, which form the most superficial layer of the skin, have begun to identify mechanisms underlying microridge morphogenesis. Distinct from other protrusions that emerge and extend as unitary structures, microridges form from the coalescence of finger-like precursor protrusions called pegs (Lam et al., 2015; Pinto et al., 2019; van Loon et al., 2020). Microridge development requires specification of apical-basal cell polarity (Magre et al., 2019; Raman et al., 2016), activity of the branched actin nucleation complex Arp2/3 (Lam et al., 2015; Pinto et al., 2019; van Loon et al., 2020), Plakin cytolinkers (Inaba et al., 2020), keratin filaments (Inaba et al., 2020), and cortical non-muscle myosin II (NMII) contraction, which concomitantly promotes apical constriction (Lam et al., 2015; Pinto et al., 2019; van Loon et al., 2020). Like microvilli, actin filaments within microridges constantly turn over (Lam et al., 2015), but the recruitment of keratin filaments by Plakin cytolinkers helps preserve microridge structure in the face of actin turnover (Inaba et al., 2020). Microridges exhibit unusual dynamics, undergoing fission and fusion to form new patterns (Lam et al., 2015). How microridge patterns mature after their initial formation has not been determined, and the molecular mechanisms executing fission and fusion are unknown.

The membranes of epithelial cells associate with a thin actomyosin filament network, called the cortex (Kelkar et al., 2020). NMII forms bipolar minifilaments within the cortex, which contract actin filaments to generate forces that regulate membrane tension, cytokinesis, and cellular morphogenesis (Kelkar et al., 2020; Martin and Goldstein, 2014). Both the density and specific arrangement of NMII minifilaments influence cortical contractility (Kelkar et al., 2020). The cortical network is attached to cell junctions, and pulls them to constrict apical surfaces during a variety of morphogenetic events (Martin and Goldstein, 2014). Cortical contraction also regulates protrusion morphogenesis. For example, contraction stimulates actin treadmilling to regulate microvillar length (Chinowsky et al., 2020). In zebrafish periderm cells, pulsatile NMII activity lowers apical membrane tension to permit the formation and elongation of microridges from peg precursors (van Loon et al., 2020). Cortical NMII contraction continues in these cells after microridges have formed (van Loon et al., 2020), but the functional significance of these later contractile events is unknown.

In this study, we characterized microridge dynamics and patterning as cells matured, and investigated the role of NMII in these processes. We found that after initial development, fission and fusion continuously remodel microridges, but these events dampen as development proceeds. High-resolution imaging revealed that cortical NMII minifilaments connect adjacent microridges, and that their specific orientation relative to microridges dictates the nature of rearrangements. These findings demonstrate that cortical NMII minifilaments are not only required for microridge formation, but also regulate microridge fission, fusion, and alignment to pattern maturing epithelial cell surfaces.

## RESULTS

### Microridge patterns mature in larval zebrafish

To determine how microridge spacing and patterning change as the developing zebrafish skin matures, we imaged zebrafish periderm cells expressing the F-actin reporter Lifeact-GFP (Riedl et al., 2008) in 48, 72 and 96 hours post-fertilization (hpf) fish (Fig 1A). Microridges had already formed and elongated by 48 hpf, but became longer on average during this period (Fig S1A-B), likely reflecting a specific reduction in pegs and short microridges (Fig S1C). Total microridge density on the apical surface increased between 48 and 96hpf (Fig 1B), which could result from an increase in microridges or reduced apical area. However, apical cell areas were not reduced, but were in fact slightly larger at 96hpf than at 48 or 72hpf (Fig S1D). Since microridge development occurs in tandem with apical constriction during early development (van Loon et al., 2020), these observations suggest that changes to microridges after 48hpf represent a distinct maturation process.

**Figure 1.**
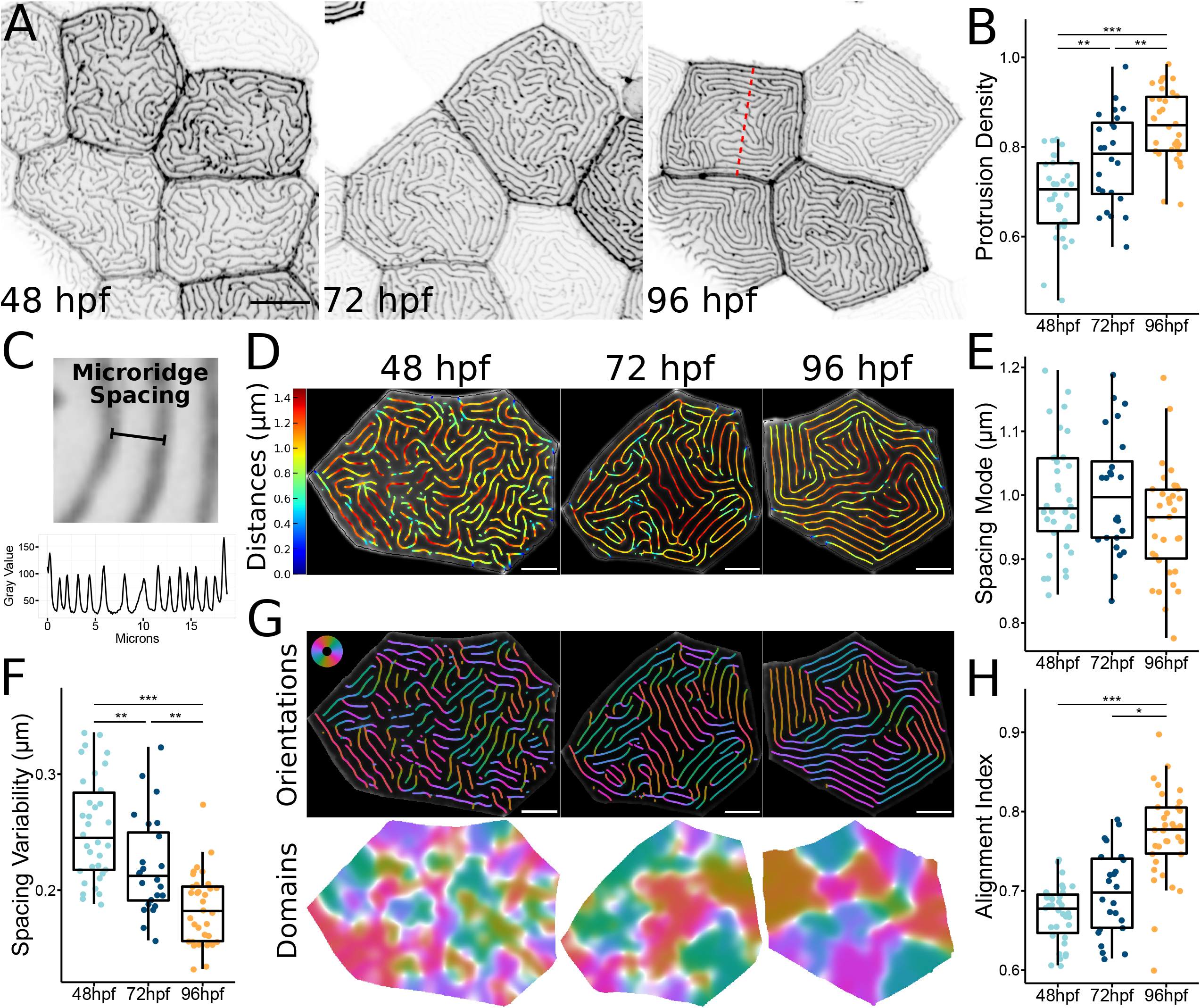
Microridge patterns mature over time. A) Representative images of periderm cells expressing Lifeact-GFP in zebrafish larvae at the specified developmental stage. Images were inverted, so that high intensity fluorescence appears black and low intensity is white. B) Dot and box-and-whisker plot of microridge density, defined as the sum microridge length (μm) normalized to apical cell area (μm^2^), on periderm cells at the specified stage. 48 hpf, n=34 cells from 12 fish; 72 hpf, n=24 cells from 10 fish; 96 hpf, n=34 cells from 15 fish. P=1.87×10^−9^, one-way ANOVA followed by Tukey’s HSD test: 48-72 hpf, P=3.32×10^−3^; 48-96 hpf, P=1.17×10^−9^; 72-96 hpf, P=6.65×10^−3^. C) Top: diagram showing definition of microridge spacing distance. Microridge spacing was measured from a given point on a microridge to the nearest point on an adjacent microridge. Bottom: profile plot of pixel intensity values across the width of a periderm cell expressing Lifeact-GFP on a 96 hpf zebrafish (indicated by dashed red line in A). D) Visualization of microridge spacing in cells at the specified developmental stage. The distance to the nearest neighboring microridge is color-coded along microridges in each cell. Colors correspond to specific distances, as indicated on the bar to the left. E) Dot and box-and-whisker plots of the mode distance between neighboring microridges in periderm cells at the specified stage. 48 hpf, n=34 cells from 12 fish; 72 hpf, n=24 cells from 10 fish; 96 hpf, n=34 cells from 15 fish. P=0.089, one-way ANOVA. F) Dot and box-and-whisker plot of microridge spacing variability, defined as the interquartile range of distances, between neighboring microridges in periderm cells at the specified stage. 48 hpf, n=34 cells from 12 fish; 72 hpf, n=24 cells from, 10 fish; 96 hpf n=34 cells from 15 fish. P=9.91×10^−10^, one-way ANOVA followed by Tukey’s HSD test: 48-72 hpf, P=7.72×10^−3^; 48-96 hpf, P=8.11×10^−10^; 72-96 hpf, P=1.90×10^−3^. G) Visualization of microridge orientation at the specified stages. Microridge orientations are color-coded along each microridge (top). Colors correspond to the color wheel on the upper left. Microridge alignment domains were expanded from microridge orientations (bottom), using the same color wheel and scale as the above microridge orientations. See Methods for details. H) Dot and box-and-whisker plot of the microridge alignment index for periderm cells at the specified stage. 48 hpf, n=34 cells from 12 fish; 72 hpf, n=24 cells from 10 fish; 96 hpf, n=34 cells from 15 fish. P=2.96×10^−12^, one-way ANOVA followed by Tukey’s HSD test: 48-72 hpf, P=0.121; 48-96 hpf, P=4.02×10^−10^; 72-96 hpf, P=8.79×10^−7^. Scale bars: 10 μm (A) and 5 μm (D and G). ‘*’ p ≥ 0.05, ‘**’ p ≥ 0.01, and ‘***’ p ≥ 0.001. For box-and-whisker plots, middle box line is the median, and lower and upper ends of boxes are 25th and 75th percentiles, respectively.

One of the most striking features of microridges is their regularly spaced and aligned arrangement, reminiscent of the parallel organization of molecules in liquid crystals, referred to as a “nematic” organization (Needleman and Dogic, 2017). To investigate how microridge spacing changes as cells mature, we measured the distance between every point on each microridge and the nearest point on a neighboring microridge (Fig 1C-D). The mode, median, and mean distances between microridges were similar between the three different stages (Fig 1E, S1E-F) and, as expected, corresponded to the orthogonal distance between aligned microridges (Fig 1C). However, spacing variability decreased over time (Fig 1F), suggesting that initially variable microridge spacing matured towards a specific spacing distance.

To determine how microridge alignment changes as microridge spacing becomes less variable, we color coding regions of cells containing microridges aligned in the same orientation. This analysis revealed that the number of domains with aligned microridges decreased, and each domain increased in area, over time (Fig 1G-H). These observations demonstrate that microridges increasingly align parallel to one another as the skin develops.

To determine if population-level changes in microridge patterning reflect microridge maturation in individual cells, we scatter-labeled periderm cells with RFP, enabling us to identify the same cells day-to-day, and thus track how microridge spacing and alignment change over time. Although each cell behaved differently, on average, microridge density increased, spacing became less variable, and microridges increasingly aligned between 48 and 96hpf (Fig 2), demonstrating that population-level trends in microridge arrangement reflect the maturation of microridge patterning in individual cells towards a nematic pattern.

**Figure 2.**
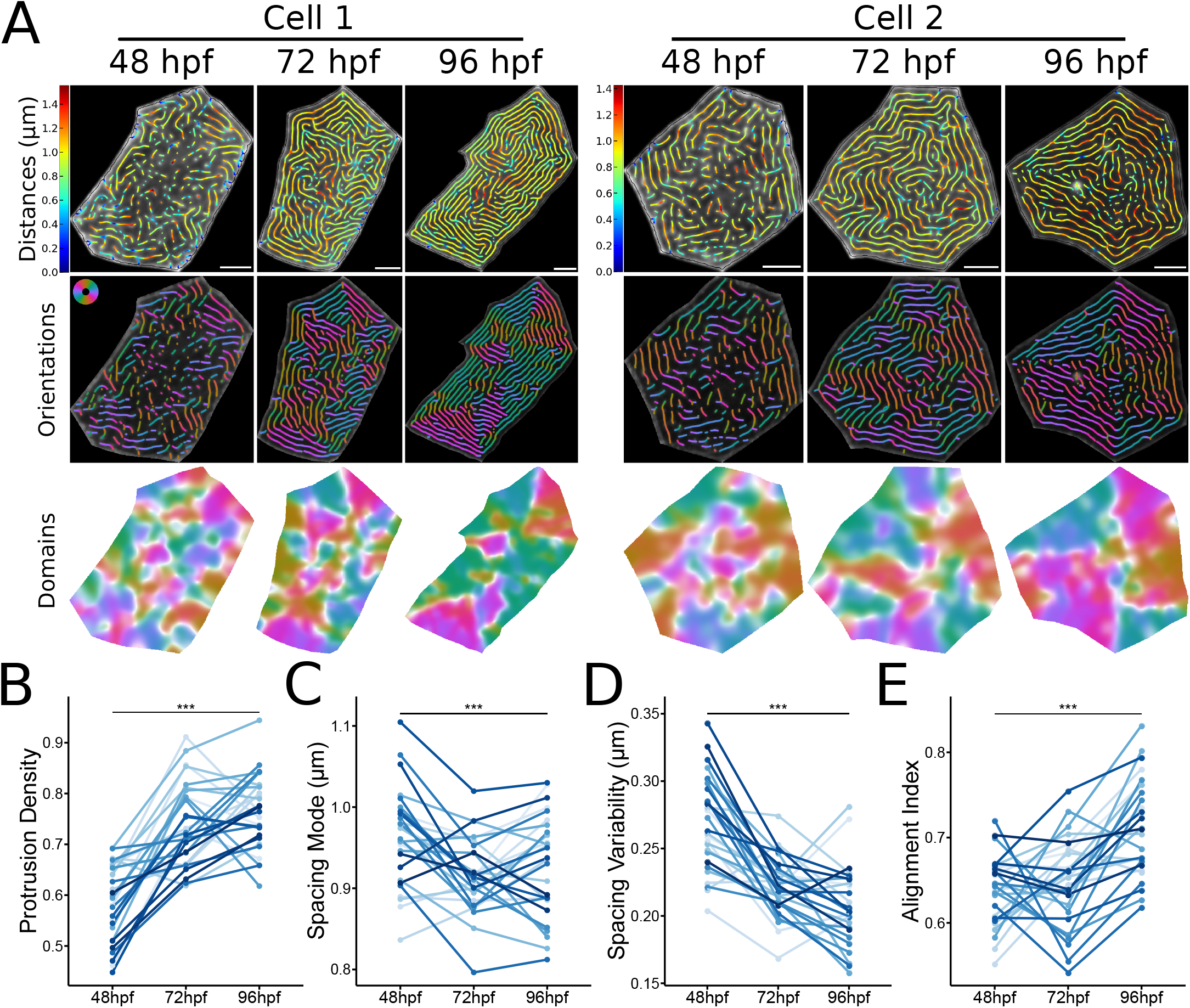
Microridge patterns mature on individual periderm cells. A) Microridge distances, orientations, and alignment domains in two cells from 48-96 hpf. B) Line and point plot of microridge density, defined as the sum microridge length (μm) normalized to apical cell area (μm^2^), in periderm cells over time. n=28 cells from 9 fish. P=5.87×10^−15^, one-way repeated measures ANOVA. C) Line and point plot of the mode distances between neighboring microridges in periderm cells over time. n=28 cells from 9 fish. P=4.08×10^−4^, one-way repeated measures ANOVA. D) Line and point plot of microridge spacing variability (interquartile range of distances) between neighboring microridges in periderm cells over time. n=28 cells from 9 fish. P=7.37×10^−11^, one-way repeated measures ANOVA. E) Line and point plot of microridge alignment index in periderm cells over time. n=28 cells from 9 fish. P=5.16×10^−8^, one-way repeated measures ANOVA. Scale bars: 5 μm (A). ‘***’ p ≥ 0.001.

### Microridges continuously rearrange

To determine the mechanism by which microridge patterns change over time, we performed time-lapse imaging of periderm cells expressing Lifeact-GFP at 30-second intervals. At each developmental stage, pegs, the finger-like precursor protrusions that coalesce to form microridges, continued to dynamically appear within and between microridges (Video 1), likely contributing to microridge lengthening. As previously observed (Lam et al., 2015), microridges underwent two types of rearrangements that altered their pattern. First, intact microridges sometimes broke apart into two separate microridges; second, two separate microridges sometimes fused end-to-end to form a longer microridge (Fig 3A, Video 1 and 2). Imaging a reporter for the plasma membrane demonstrated that these events reflect fission or fusion of the whole protrusion, not just of its internal actin structure (Figure 3B, Video 3). As microridges matured, rearrangement events decreased from 0.362 events/μm · min at 48 hpf to 0.155 and 0.115 events/μm · min at 72 and 96 hpf, respectively (Fig 3C, Video 1). Fission and fusion events occurred with roughly equal frequency, and this proportion did not change over time (Fig 3D), but the frequency of these rearrangements decreased as the pattern matured (Fig 3E, Spearman Correlation Coefficient = −0.832).

**Figure 3.**
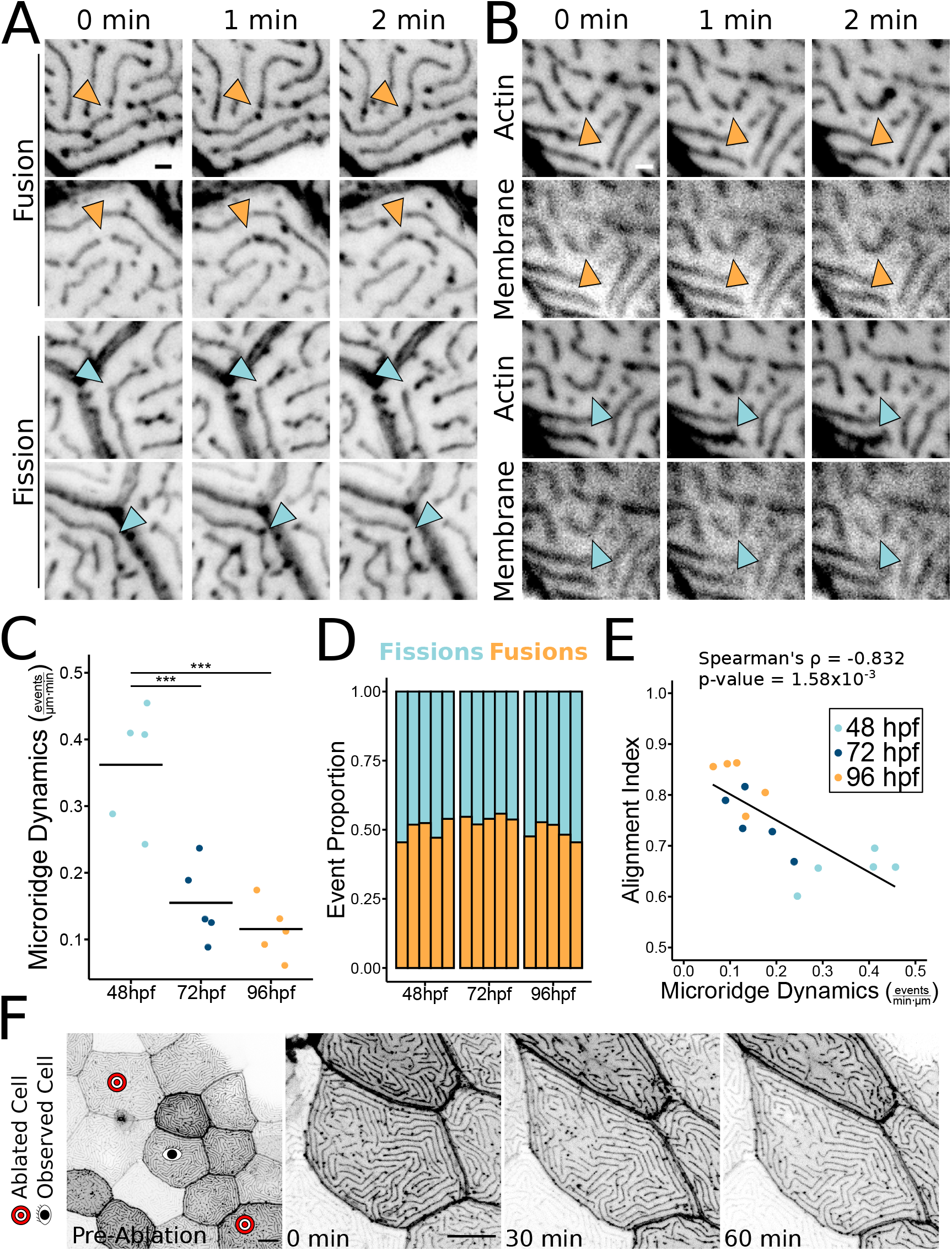
Microridges dynamically rearrange. A) Time-lapse images of 48 hpf zebrafish periderm cells expressing Lifeact-GFP demonstrating microridges undergoing fission or fusion. Orange arrowheads indicate fusion; blue arrowheads indicate fission. Images were inverted, so that high intensity fluorescence appears black and low intensity is white. Images are still frames from Video 2. B) Time-lapse images of 48hpf zebrafish periderm cells expressing Lifeact-GFP (actin) and mRuby-PH-PLC (membrane) demonstrating microridges undergoing fission or fusion. Orange arrowheads indicate fusion; blue arrowheads indicate fission. Images were inverted, so that high intensity fluorescence appears black and low intensity is white. Images are still frames from Video 3. C) Jittered dot plot of the sum of fission and fusion events in each cell, normalized to cell apical area, over a 10-minute period (events/μm · min) at the specified stage. Middle bar represents the mean. n=5 cells from 5 fish at all stages. P=1.62×10^−4^, one-way ANOVA followed by Tukey’s HSD test: 48-72 hpf, P=9.75×10^−4^; 48-96 hpf, P=2.17×10^−4^; 72-96 hpf, P=0.629. D) Stacked bar plot of the proportion of fission and fusion events indicated stages. Each bar represents one cell. n=5 cells from 5 fish at all developmental stages. Fusion events were used in a Test of Equal Proportions (P=0.224) and fusion event proportion estimates were 0.501, 0.543, and 0.488 for 48, 72, and 96hpf, respectively. E) Scatter plot of microridge dynamics (events/μm · min) over a 10-minute period versus the microridge alignment index at the start of the 10-minute period. n=5 cells from 5 fish at all developmental stages. P=1.16×10^−4^, Spearman’s rank correlation rho = −0.832. F) Time-lapse image sequence of Lifeact-GFP-expressing periderm cells stretching in response to neighbor cell ablation on a 72 hpf zebrafish. Pre-ablation image shows the cell of interest between the ablated cells, and images of the cell of interest immediately after ablation (0 min) and at 30-minute intervals after ablation. The cell stretched dramatically, but microridge rearrangements did not appreciably increase. Images were inverted, so that high intensity fluorescence appears black and low intensity is white. Images are still frames from Video 4. Scale bars: 1 μm (A and B) and 10 μm (E). ‘***’ p ≥ 0.001.

### Cell stretching does not induce microridge rearrangement

Periderm cells are constantly pushed and pulled by neighboring cells as the epidermis grows. We therefore speculated that microridge fission and fusion may be induced by forces associated with cell shape distortion. To test this idea, we ablated periderm cells on either side of an observed cell using a laser on a 2-photon microscope (O’Brien et al., 2009b; van Loon et al., 2020). This procedure caused the central cell to stretch between the two wounds, and often pucker or bulge in the orthogonal axis. Surprisingly, cell elongation did not increase microridge fission or fusion, but simply distorted microridges to accommodate the cells’ new shapes (Fig 3F, Video 4). This observation suggests that microridges do not undergo fission or fusion simply as a result of cellular distortion, and thus implies that remodeling events are actively regulated.

### Microridge rearrangements require cortical NMII contraction

The apical cortex of periderm cells experiences pulsatile NMII-based contractions through at least 48 hpf (van Loon et al., 2020). These contractions are required for apical constriction and the coalescence of peg precursors to form and elongate microridges (van Loon et al., 2020), but later functions have not been described. To test if cortical contraction affects microridge fission or fusion events, we made time-lapse videos of periderm cells expressing fluorescent reporters for both F-actin (Lifeact-Ruby) and NMII (Myl12.1-EGFP) (Maître et al., 2012; van Loon et al., 2020). At 48hpf, periderm cells displayed local pulses of NMII reporter fluorescence in the apical cortex (Fig 4A, Video 5), which we previously found to reflect NMII contraction (van Loon et al., 2020). Many of these contraction events correlated spatially and temporally with microridge rearrangements (Fig 4A). To quantify this correlation, we measured the distance between microridge rearrangement events and the nearest detectable NMII contractile pulse in the same frame. On average, 41% of microridge rearrangements occurred within 1μm of an NMII contraction (Fig 4B). By contrast, when the NMII reporter channel was rotated 90°, only 22% occurred within 1μm of a contraction (Fig 4B), indicating that the coincidence between contraction and rearrangement events did not occur by chance. These observations likely underestimate the number of rearrangement events associated with contraction, since contractions may be shorter-lasting or dimmer than we can detect with our reporter. NMII contractions equally correlated with fission and fusion events (Fig 4C).

**Figure 4.**
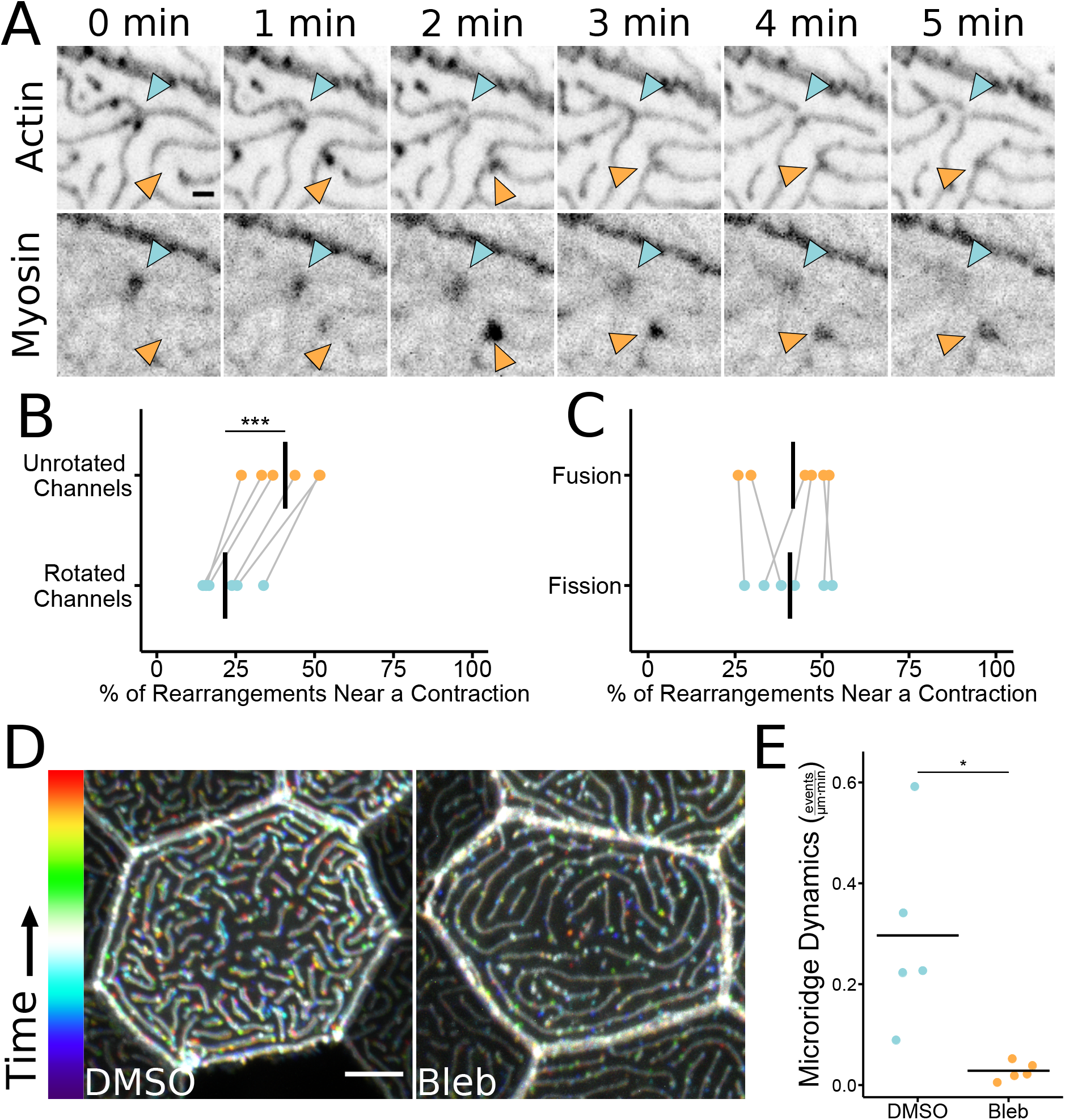
Microridge rearrangements spatially and temporally correlate with NMII contraction. A) Time-lapse images of 48 hpf zebrafish periderm cells expressing Lifeact-mRuby (actin) and Myl12.1-EGFP (myosin). Microridges fission as a myosin contraction dissipates (blue arrowheads). Microridges fuse as a myosin contraction intensifies (orange arrowheads). Images were inverted, so that high intensity fluorescence appears black and low intensity is white. Images are still frames from Video 5. B) Dot plot of the percentage of microridge fission and fusion events within 1μm of an NMII contraction over a 10-minute period. Graph compares unrotated channels to data analyzed after rotating the NMII fluorescence channel 90° relative to the actin fluorescence channel. Grey lines connect the unrotated samples to their rotated counterparts. n=6 cells from 6 fish, including 3 cells from 3 fish at 24 hpf and 3 cells from 3 fish at 48 hpf. P=2.27×10^−4^, paired t-test. C) Dot plot of the percentage of microridge fission and fusion events within 1 μm of an NMII contraction over a 10-minute period. Graph compares contraction-associated fusion events to contraction-associated fission events in the same cells. Grey lines connect points from the same cell. n=6 cells from 6 fish, including 3 cells from 3 fish at 24 hpf and 3 cells from 3 fish at 48 hpf. P=0.778, paired t-test. D) Representative color-coded temporal projections (corresponding to the bar on the left) of images from 10-minute time-lapse movies (30-second intervals) in 49 hpf zebrafish periderm cells expressing Lifeact-GFP after 1hr exposure to 1% DMSO (vehicle control) or 50μM blebbistatin. Microridges in control cells are more dynamic over the 10-minute period, so control cells appear more colorful than blebbistatin-treated cells. Images are temporal projections of Video 6. E) Jittered dot plot of the sum of fission and fusion events in each cell, normalized to cell apical area, over a 10-minute period (events/μm · min) in cells after 1-hour exposure to 1% DMSO (vehicle control) or 50μM blebbistatin. n=5 cells from 5 fish for control and treatment. P=0.033, unpaired t-test. Scale bars: 5μm (D). ‘*’ p ≥ 0.05 and ‘***’ p ≥ 0.001. Bars in dot plots represent the mean.

To directly test if NMII contraction is required for microridge rearrangements, we treated 48hpf fish with the specific NMII inhibitor blebbistatin (Straight et al., 2003) for one hour, then made 10-minute videos of periderm cells expressing Lifeact-GFP. NMII inhibition dramatically reduced fission and fusion compared to controls (Fig 4D-E, Video 6), demonstrating that NMII activity is required for microridge remodeling.

### NMII contraction regulates microridge density and spacing

Given that NMII contraction promotes microridge rearrangements, and that these dynamic events negatively correlate with microridge alignment, we hypothesized that inhibiting NMII contraction may disrupt microridge maturation. To determine the long-term consequences of suppressing NMII activity, we treated zebrafish with blebbistatin for 24 hours, starting at 48hpf. Compared to controls, microridges in blebbistatin-treated animals were shorter, distributed less densely, and spaced more widely (Fig 5A-E). These observations indicate that microridges must be actively maintained by contraction, which may facilitate the incorporation of new pegs into established microridges. Blebbistatin also increased microridge alignment. This effect on alignment may be a consequence of the lower microridge density, since our alignment index measures the number of domains containing aligned microridges (Fig 5B,D,F), but could also indicate that suppressing contraction allows the system to settle into a local energy minimum (see Discussion). Since long-term NMII inhibition can have deleterious, indirect effects on cells, we compared microridges on individual cells before and after 1-hour blebbistatin treatment. Similar to 24-hour treatment, 1-hour exposure to blebbistatin disrupted microridge spacing, decreasing density and increasing the microridge alignment index (Fig 6).

**Figure 5.**
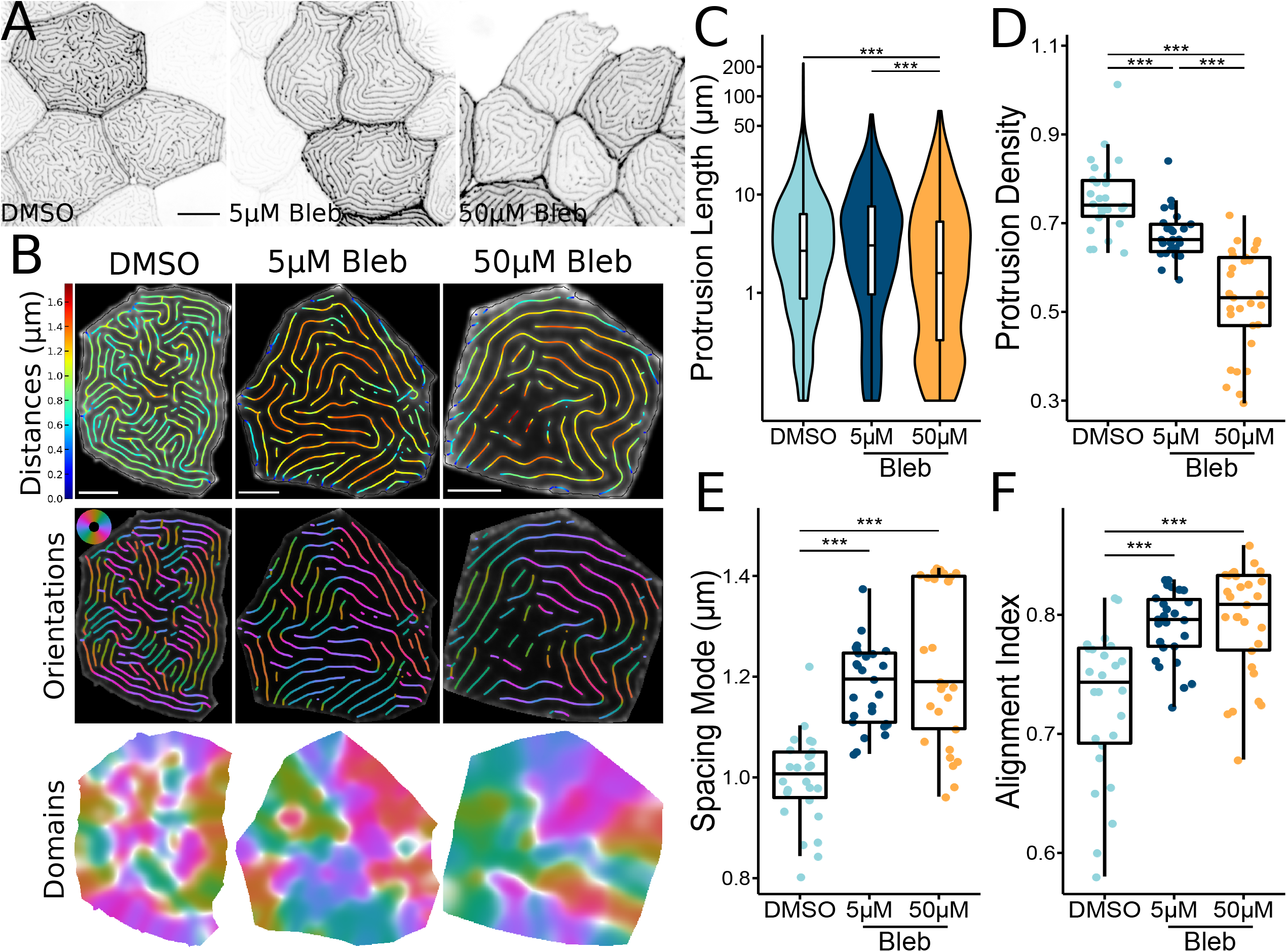
Inhibiting NMII changes microridge patterns. A) Representative images of periderm cells expressing Lifeact-GFP on 72 hpf zebrafish after 24-hour exposure to the specified concentration of blebbistatin or vehicle control (DMSO). Images were inverted, so that high intensity fluorescence appears black and low intensity is white. B) Visualizations of microridge distances, orientations, and alignment domains from periderm cells at 72 hpf after 24-hour exposure to the specified concentration of blebbistatin or vehicle control (DMSO). C) Violin and box-and-whisker plot of projection length for periderm cells in 72 hpf zebrafish after 24-hour exposure to the specified concentration of blebbistatin or vehicle control (DMSO). DMSO, n=26 cells from 9 fish; 5μM blebbistatin, n=27 cells from 9 fish; 50μM blebbistatin, n=29 cells from 9 fish. P<2.2×10^−16^, Kruskal-Wallis test followed by Dunn test with Benjamini-Hochberg p-value adjustment: DMSO-5μM blebbistatin, P=0.173; DMSO-50μM blebbistatin, P=2.51×10^−13^; 5μM blebbistatin-50μM blebbistatin, P=3.79×10^−17^. D) Dot and box-and-whisker plot of microridge density, defined as the sum microridge length (μm) normalized to apical cell area (μm^2^), for periderm cells in 72 hpf zebrafish after 24-hour exposure to the specified concentration of blebbistatin or vehicle control (DMSO). DMSO, n=26 cells from 9 fish; 5μM blebbistatin, n=27 cells from 9 fish; 50μM blebbistatin, n=29 cells from 9 fish. P=2.80×10^−14^, one-way ANOVA followed by Tukey’s HSD test: DMSO-5μM blebbistatin, P=3.07×10^−3^; DMSO-50μM blebbistatin, P<2×10^−16^; 5μM blebbistatin-50μM blebbistatin, P=7.70×10^−8^. E) Dot and box-and-whisker plot of the mode distance between neighboring microridges in periderm cells in 72 hpf zebrafish after 24-hour exposure to the specified concentration of blebbistatin or vehicle control (DMSO). DMSO, n=26 cells from 9 fish; 5μM blebbistatin, n=27 cells from 9 fish; 50μM blebbistatin, n=29 cells from 9 fish. P=0.318, one-way ANOVA. F) Dot and box-and-whisker plot of the alignment index on periderm cells in 72 hpf zebrafish after 24-hour exposure to the specified concentration of blebbistatin or vehicle control (DMSO). DMSO, n=26 cells from 9 fish; 5μM blebbistatin, n=27 cells from 9 fish; 50μM blebbistatin, n=29 cells from 9 fish. P=4.56×10^−7^, one-way ANOVA followed by Tukey’s HSD test: DMSO-5μM blebbistatin, P=1.11×10^−5^; DMSO-50μM blebbistatin, P=2.38×10^−6^; 5μM blebbistatin-50μM blebbistatin, P=0.951. Scale bars: 10uM (A) and 5μm (B). ‘**’ p ≥ 0.01 and ‘***’ p ≥ 0.001. For box-and-whisker plots, middle box line is the median, and lower and upper ends of boxes are 25th and 75th percentiles, respectively.

**Figure 6.**
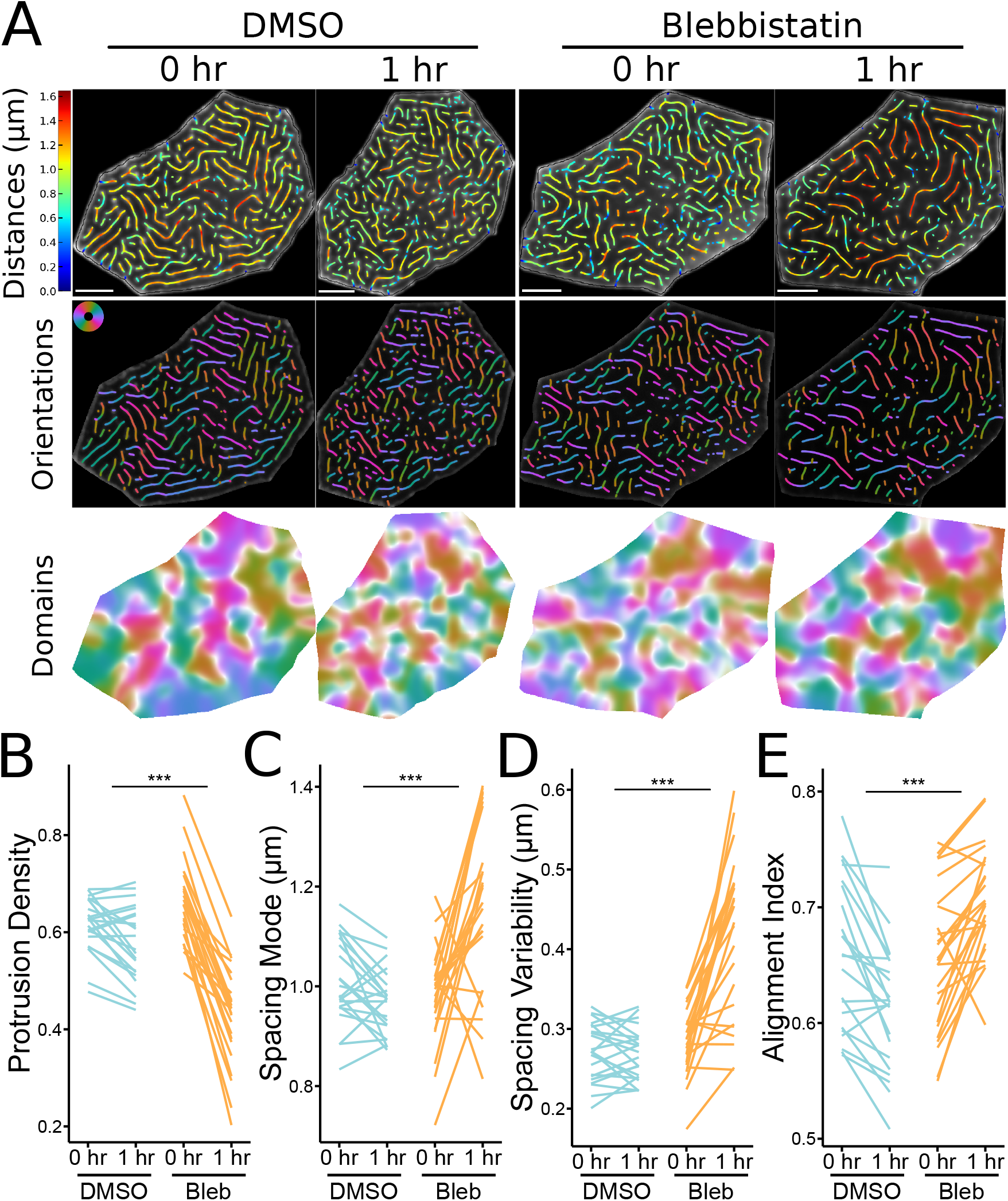
Short-term inhibition of NMII contractility alters microridge patterns in individual cells. A) Representative visualizations of microridge distances, orientations, and alignment domains in periderm cells expressing Lifeact-GFP before (48 hpf, 0 hr) and after (49 hpf, 1hr) 1-hour treatment with 50μM blebbistatin or vehicle (DMSO). B) Line plot of microridge density, defined as the sum microridge length (μm) normalized to apical cell area (μm^2^), from periderm cells before (48 hpf, 0 hr) and after (49 hpf, 1hr) 1-hour treatment with 50μM blebbistatin or vehicle control (DMSO). DMSO, n=22 cells from 4 fish; 50μM blebbistatin, n=25 cells from 4 fish. P=4.09×10^−11^, one-way repeated measures ANOVA. C) Line plot of microridge spacing mode from periderm cells before (48 hpf, 0 hr) and after (49 hpf, 1hr) 1-hour treatment with 50μM blebbistatin or vehicle control (DMSO). DMSO, n=22 cells from 4 fish; 50μM blebbistatin, n=25 cells from 4 fish. P=7.76×10^−6^, one-way repeated measures ANOVA. D) Line plot of microridge spacing variability (interquartile range of distances) between neighboring microridges in periderm cells before (48 hpf, 0 hr) and after (49 hpf, 1hr) 1-hour treatment with 50μM blebbistatin or vehicle control (DMSO). DMSO, n=22 cells from 4 fish; 50μM blebbistatin, n=25 cells from 4 fish. P<2×10^−16^, one-way repeated measures ANOVA. E) Line plot of the alignment index in periderm cells before (48 hpf, 0 hr) and after (49 hpf, 1hr) 1-hour treatment with 50μM blebbistatin or vehicle control (DMSO). DMSO, n=22 cells from 4 fish; 50μM blebbistatin, n=25 cells from 4 fish. P=1.02×10^−8^, one-way repeated measures ANOVA. Scale bars: 5μm (A). ‘***’ p ≥ 0.001.

### High-resolution imaging reveals individual NMII minifilaments in the cortex

Since NMII inhibition experiments could not disambiguate NMII’s role in regulating microridge fission and fusion, length maintenance, and spacing, we addressed these questions by imaging NMII organization and activity in the periderm cortex directly. To image NMII and F-actin with improved spatial resolution, we used Airyscan microscopy (Weisshart, 2014). Using this approach, the NMII reporter often appeared as pairs of puncta (Fig 7A). The Myl12.1-EGFP NMII reporter is a fusion of EGFP to a myosin regulatory light chain (Maître et al., 2012; van Loon et al., 2020), which binds near myosin heads at opposing ends of NMII minifilaments. We thus speculated that puncta pairs represent ends of single bipolar minifilaments. Consistent with this possibility, the median distance between intensity maxima of NMII reporter doublets was 281nm (Fig 7B), similar to the reported length of bipolar minifilaments assembled in vitro (~300nm in length; (Billington et al., 2013)). To further test if these structures are individual minifilaments, we imaged periderm cells expressing reporters for both NMII light chain (Myl12.1-Ruby) and a C-terminally tagged NMII heavy chain (Myh9a-EGFP). A fluorophore at the C-terminus of NMII heavy chains should localize to the middle of minifilaments, between NMII heavy chain heads (Fig 7C-D). Puncta in periderm cells expressing both reporters were arranged in the expected alternating pattern (Fig 7C-D). Thus, our imaging system allows us to distinguish individual NMII minifilaments within the plane of the apical cortex in cells of living animals.

**Figure 7.**
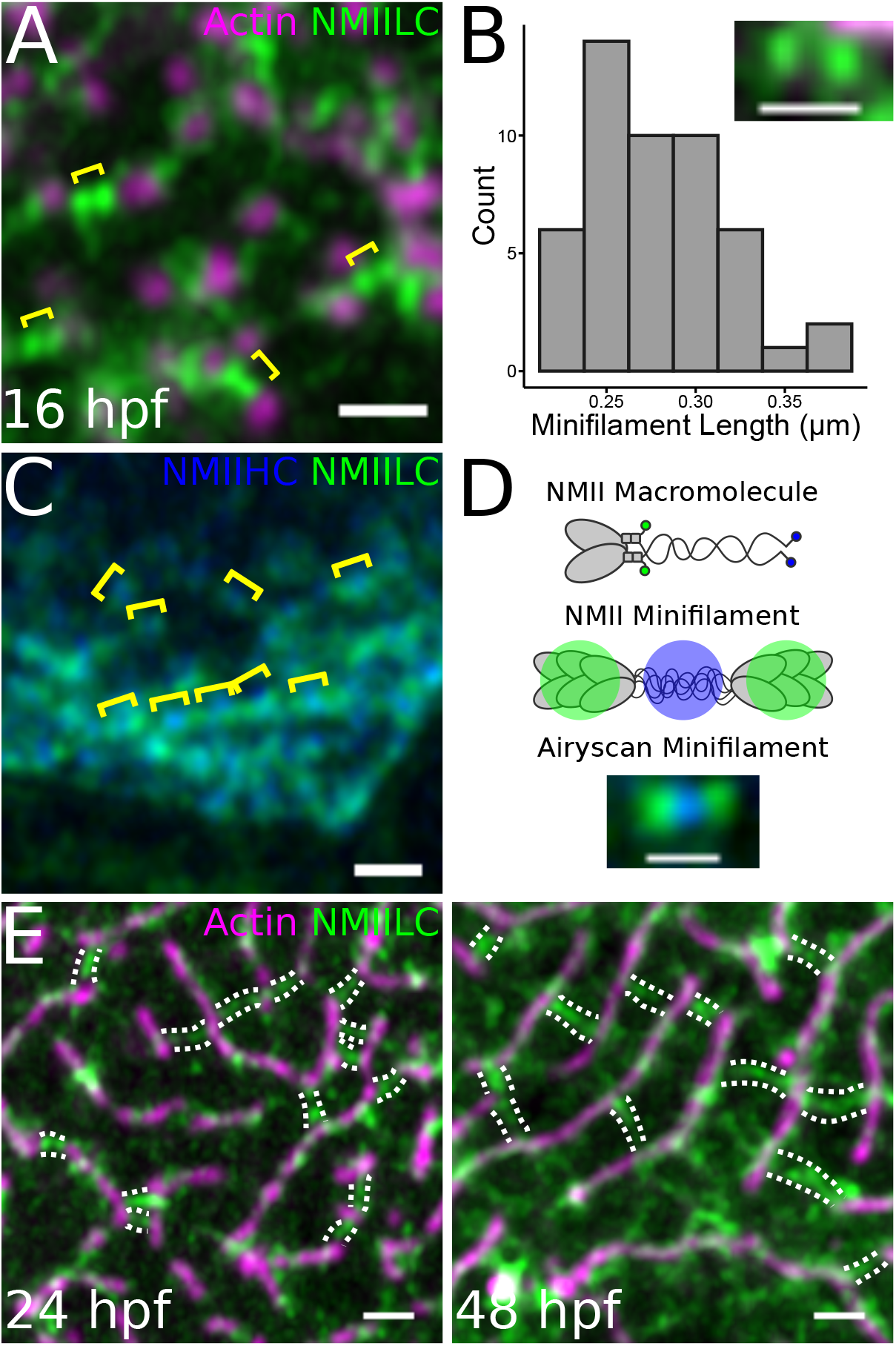
NMII minifilaments connect adjacent pegs and microridges. A) Airyscan image of a 16 hpf zebrafish periderm cell expressing fluorescent reporters for actin (Lifeact-Ruby) and NMII light chain (Myl12.1-GFP). Pairs of green puncta (yellow brackets) appear in the cortex between adjacent pegs (magenta puncta). B) Histogram of distances between the intensity maxima of presumptive NMII minifilaments. Inset is a representative image showing GFP signal at opposing ends of a presumptive NMII minifilament in a periderm cell expressing reporters for actin (Lifeact-Ruby) and NMII light chain (Myl12.1-GFP). n=49 minifilaments from 4 cells on 4 fish. C) Airyscan image of a 24 hpf zebrafish periderm cell expressing fluorescent reporters for NMII heavy chain (NMIIHC, Myh9a-mCherry) and NMII light chain (NMIILC, Myl12.1-GFP). NMIIHC channel was pseudo-colored blue. Brackets show examples of GFP-mCherry-GFP fluorescence patterns. D) Diagram of NMII fluorescent protein fusion design and expected NMII minifilament fluorescence pattern. The upper graphic shows an NMII macromolecule, composed of two heavy chains, two essential light chains, and two regulatory light chains. GFP was fused to the regulatory light chains (Myl12.1-GFP), while mCherry was fused to the tail of the heavy chains (Myh9a-mCherry; represented in blue). The middle graphic shows the expected fluorescence pattern when multiple NMII macromolecules, labelled like the one in the upper graphic, assemble into an NMII minifilament. The lower Airyscan image shows an NMII minifilament in the cortex of a 24hpf zebrafish periderm cell expressing Myl12.1-GFP and Myh9a-mCherry. E) Airyscan images showing NMII minifilaments connecting adjacent microridges side-to-side and end-to-end during (24hpf) and after (48hpf) microridge formation in periderm cells expressing reporters for actin (Lifeact-Ruby) and NMII light chain (Myl12.1-GFP). Dotted lines track along NMII minifilament “bridges”. Scale bars: 1 μm (A, C, & E) and 500 nm (B & D)

### Cortical NMII minifilaments associate with pegs and microridges

To determine how NMII minifilaments are arranged relative to cell protrusions, we imaged them, along with F-actin, at several developmental stages. Prior to microridge formation (16 hpf), NMII minifilaments in the apical cortex were closely associated with microridge peg precursors (Fig 7A), and continued to associate with protrusions as pegs coalesced to form microridges. At 24 hpf, NMII minifilaments were often attached to two separate microridges, bridging them end-to-end or side-to-side (Fig 7E). This organization was maintained as microridges matured: At 48hpf many cortical NMII “bridges” formed perpendicular connections between adjacent microridges, often appearing to consist of two end-to-end minifilaments (Fig 7E).

### NMII minifilaments orchestrate microridge rearrangement and spacing

To observe how the organization of NMII minifilaments in the cortex relates to protrusion dynamics, we made high-resolution videos of periderm cells expressing Lifeact-Ruby and Myl12.1-EGFP. During early morphogenesis, appearance and disappearance of pegs often correlated with appearance and disappearance of NMII reporter signal, and movement of pegs was associated with a corresponding movement of the NMII reporter (Fig 8A, Video 7), confirming that NMII minifilaments are tethered to protrusions. At later stages, when microridges remodel, the orientation of NMII minifilaments correlated with the type of microridge rearrangement observed. Minifilaments connecting the ends of two microridges appeared to pull them together, fusing them into a longer microridge (Fig 8B, Video 7). By contrast, minifilaments oriented perpendicular to microridges were often associated with fission events, which occurred at the point where microridges attached to the minifilaments (Fig 8B, Video 7). Minifilaments arranged perpendicular to microridges also appeared to regulate microridge spacing: the attachment of minifilaments to two parallel microridges appeared to bring them closer together, whereas their disappearance or detachment allowed the two microridges to drift apart (Fig 8C, Video 7). These observations suggest that the spatial organization of individual cortical NMII minifilaments directs the rearrangement of microridges, regulating the spacing between them and thus altering the microridge pattern on the apical surface.

**Figure 8.**
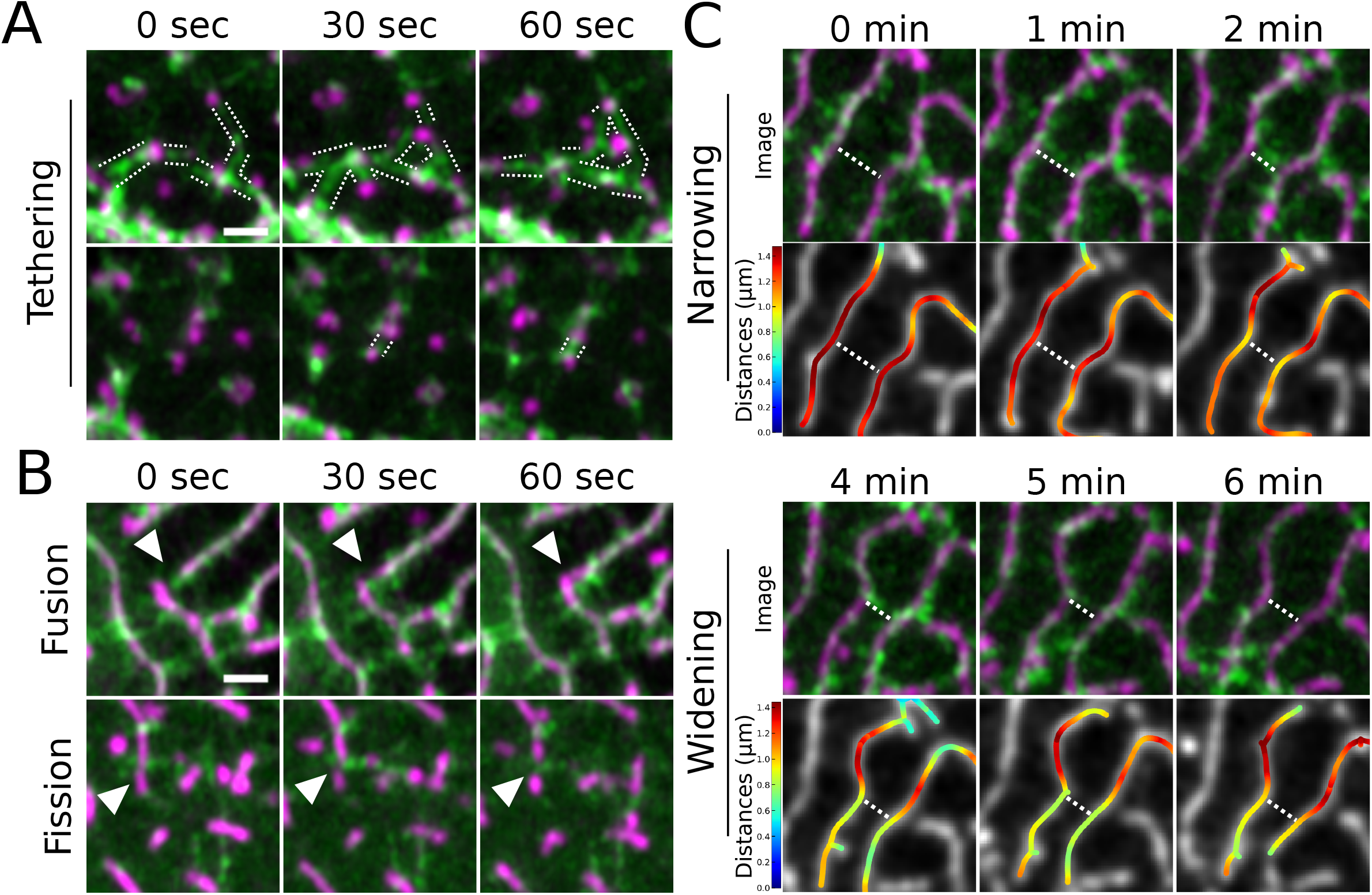
NMII minifilaments dynamically connect pegs and organize microridge rearrangements. A) Airyscan time-lapse images of NMII minifilaments dynamically connecting pegs as they emerge in the cortex of a periderm cell expressing fluorescent reporters for actin (Lifeact-mRuby) and NMII (Myl12.1-GFP). Dotted lines track along NMII minifilament “bridges”. Images include still frames from Video 7. B) Airyscan time-lapse images of microridge rearrangements (white arrowheads) in periderm cells expressing fluorescent reporters for actin (Lifeact-mRuby) and NMII (Myl12.1-GFP). In the upper panels, an NMII minifilament connects the ends of adjacent microridges, fusing them together. In the lower panels, NMII minifilaments line up perpendicular to a microridge and appear to sever it. Images are still frames from Video 7. C) Airyscan time-lapse images of microridge spacing dynamics in periderm cells expressing fluorescent reporters for actin (Lifeact-mRuby) and NMII (Myl12.1-GFP). Dotted line highlights narrowing and widening regions. Top rows show Airyscan images; bottom rows show color-coded distances. Colors correspond to color bars on the left. The upper two rows show NMII minifilaments connecting two adjacent microridges and apparently pulling them together. The lower two rows show the NMII minifilament bridge between microridges dissipating as the adjacent microridges separate. Images are still frames from Video 7. Scale bars: 1μm (A and B).

## DISCUSSION

Our study reveals that cortical NMII orchestrates a unique process for the patterning and maturation of microridges. Cells retain microridges on their surfaces for days, and likely even weeks, but, unlike extremely stable stereocilia, microridges continuously remodel through an NMII-mediated “recombination” process of fission and fusion as they mature towards a more ordered, nematic arrangement (Fig 9). Thus, at least during the first week of development, microridges are not permanent cell identifiers, like a fingerprint, but evolving structures that form new patterns over time.

**Figure 9.**
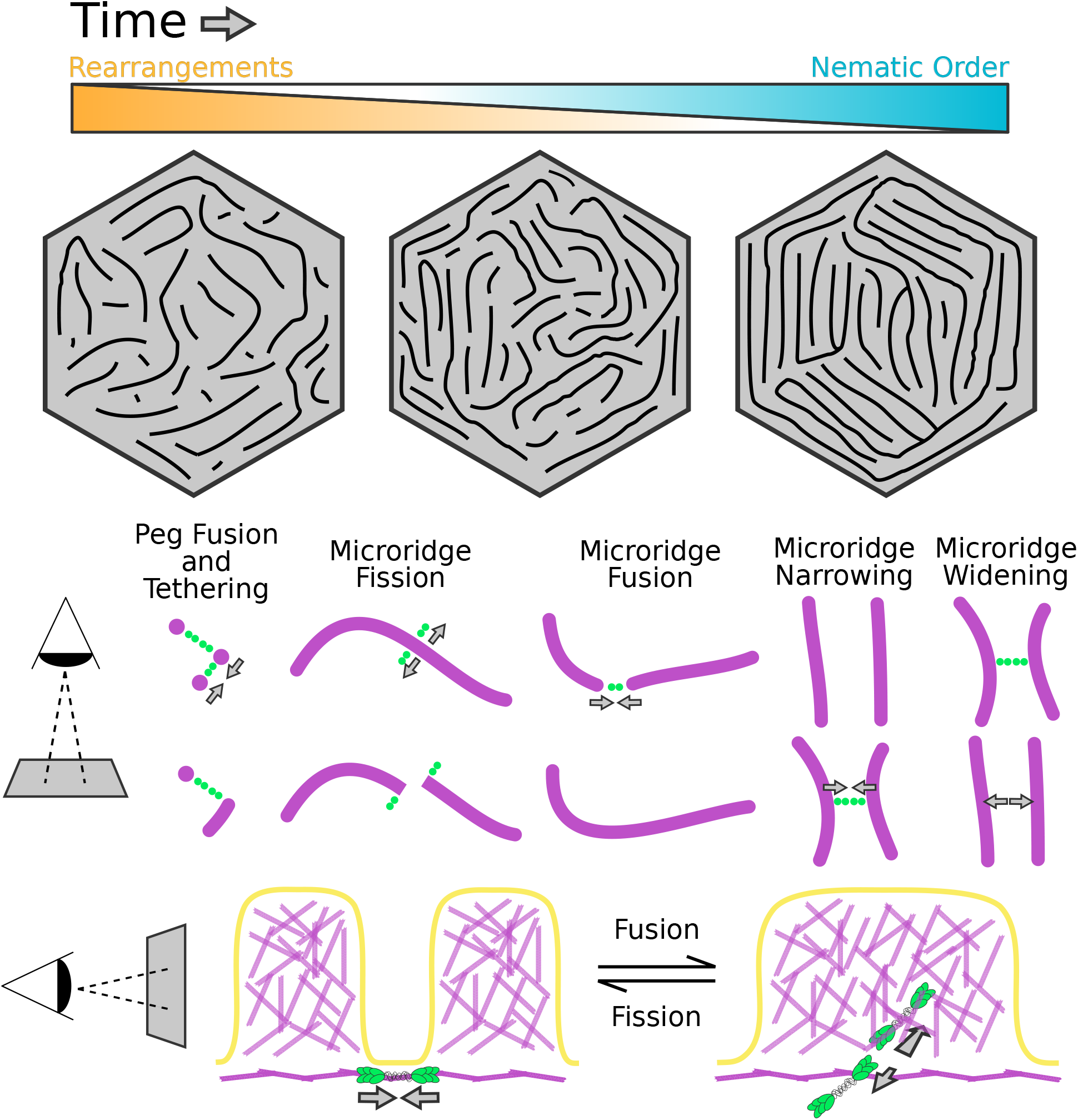
Model for microridge maturation and minifilament-mediated rearrangements. Top: The nematic order of microridge patterns increases as rearrangements decrease in frequency. Bottom: The orientation of NMII minifilaments determines the outcome of rearrangement events and regulates spacing (see Discussion).

### Microridge pattern maturation minimizes surface energy

The increasing nematic order of maturing microridge patterns suggests that they are governed by an energy minimization principle, which can be explained with concepts defined by physics. Optimal parallel packing of microridges likely minimizes the bending and stretching energy of the lipid bilayer that is coupled to the underlying cortex. Consistent with this idea, we found that the size of local alignment domains increased, and their number decreased, as microridge patterns matured (Fig 1, Fig 2). Inevitably, initial disorder in the emerging pattern brings about sharp boundaries between the domains of local alignment. These boundaries are defects in the nematic order, and thus associated with an energy penalty, a phenomenon well-known in liquid crystals (Needleman and Dogic, 2017). The global energy minimum likely corresponds to concentric microridges arranged in parallel rings, like a target. Our observations show that microridge patterns, which are initially in states with many alignment domains, progress towards this well-ordered global minimum over time, a process that requires crossing energy barriers associated with fission and fusion of preexisting microridges. However, this global minimum cannot be readily reached from an arbitrary state with multiple alignment domains without crossing energy barriers associated with fission and fusion of preexisting microridges.

Our results suggest that myosin activity facilitates the crossing of energy barriers, promoting fission and fusion and thus rearranging microridge patterns. The fact that myosin activity leads to microridge fission and fusion with approximately equal probability (Fig 3D) suggests that it does not increase their order or disorder per se, but rather provides quantal “kicks” that locally alter pattern topology. Thus, myosin activity is analogous to the thermodynamic temperature of the pattern--by randomly breaking and fusing individual microridges, myosin allows the pattern to cross energy barriers separating local energy minima. Following this thermodynamic analogy, the decrease in microridge rearrangement events over time corresponds to slowly lowering the temperature, or annealing, which is well-known in physics to help systems reach lower energy states on complex energy landscapes with multiple minima (van Laarhoven and Aarts, 1987). Blebbistatin may represent rapid quenching (a sharp temperature drop) that allows the system to descend into the closest energy minimum, perhaps explaining why blebbistatin in our experiments increased the alignment index.

### Microridges are modular protrusions

Both the initial formation and remodeling of microridges demonstrate that they are modular structures: individual units (pegs) assemble into longer structures (microridges); once assembled, microridges can be broken at any point and attached to other microridges. This modular nature distinguishes microridges from other protrusions. However, the apparent simplicity of this process elides the complexity of rearrangement events at the molecular level, which likely involve multiple, locally regulated activities. Fission requires not just severing actin filaments, but also locally disassembling a supramolecular network of F-actin, keratin filaments, and actin-binding proteins (Pinto et al., 2019), as well as membrane remodeling. Microridge remodeling events require NMII activity, but fission is likely instigated by upstream regulators that coordinate multiple biochemical activities. Such roles could be played by Rho family GTPases, which can regulate both F-actin stability and NMII contraction (Kelkar et al., 2020; Ridley, 2015), or Aurora B kinase, which promotes NMII activity (Minoshima et al., 2003; Touré et al., 2008) and disassembly of actin and keratin filaments (Field et al., 2019) at the cytokinetic furrow. Fusion likely requires F-actin polymerization, the activity of F-actin cross-linking proteins that link the cytoskeletal networks of the two parent microridges, and the reintegration of keratin filaments, which have the potential to connect with themselves end-to-end (Çolakoğlu and Brown, 2009). NMII activity may facilitate both fission and fusion by reducing surface tension, as it does during initial microridge formation (van Loon et al., 2020), and by physically pulling protrusions to rupture them or bring them together.

### The orientation of cortical NMII minifilaments determines the nature of microridge rearrangements

Visualizing individual NMII minifilaments in the cortex of living cells provided insight into how they execute microridge rearrangements, as well as evidence that they play a direct role in microridge spacing (Fig 9). From the earliest steps of microridge morphogenesis, cortical minifilaments associate with protrusions. This observation suggests that the ends of individual minifilaments are biochemically tethered to the base of pegs and microridges, orthogonal to actin filaments in these protrusions. When two pegs are tethered to opposite ends of a minifilament, contraction brings them closer together, providing an opportunity for them to fuse into a nascent microridge. Similarly, contraction of peg-to-microridge minifilament bridges may contribute to microridge elongation, and contraction of minifilament bridges connecting two microridge ends may promote microridge fusion. The recruitment of keratin filaments into these growing protrusions likely helps stabilize them (Inaba et al., 2020), preventing fusion events from reversing. By contrast, minifilaments tethered to the sides of microridges sometimes promoted fission, suggesting that minifilaments may pull on microridges to facilitate their local disassembly. If minifilaments bridged two parallel microridges, they often appeared to pull them closer together without severing them, providing direct evidence that NMII can regulate microridge spacing. At the later stages, microridges were often linked by a bridge of two perpendicular minifilaments aligned end-to-end. This arrangement raises the intriguing possibility that minifilaments could serve as molecular spacers for aligned microridges, similar to how spectrin tetramers determine the spacing of actin rings in axons (Xu et al., 2013). However, it is at least as likely that spacing length is determined by another factor, such as the minimization of membrane bending energy, and that minifilaments organize to accommodate that spacing.

Minifilaments are typically thought to be isotropically oriented in the cortex of interphase cells (Kelkar et al., 2020), but our findings suggest that their association with microridges causes them to adopt a highly organized arrangement in the cortex of periderm cells. In mature periderm cells, since microridges align with cell borders and with one another, their arrangement approximates an ideal target-like concentric pattern; because minifilaments form perpendicular bridges between adjacent microridges, they are predominantly arranged radially towards the center of cells. Since contractility is influenced not just by minifilament density, but also the relative arrangement of NMII in the cortex (Kelkar et al., 2020), this unusual radial minifilament organization likely endows periderm cells with unique contractile properties. Collectively, our observations reveal a surprisingly organized arrangement of cortical minifilaments, indicating that understanding how cortical contraction executes specific biological processes will require a better understanding of cortical minifilament architecture.

## MATERIALS AND METHODS

### Zebrafish

Zebrafish (*Danio rerio*) were raised at 28.5°C on a 14-h/10-h light/dark cycle. Embryos were raised at 28.5°C in embryo water composed of 0.3 g/Liter Instant Ocean salt (Spectrum Brands, Inc.) and 0.1% methylene blue. Previously characterized zebrafish lines in this paper include AB wild-type fish (ZFIN: ZDB-GENO-960809-7), Tg(krt5:Gal4) (Rasmussen et al., 2015), Tg(UAS:Lifeact-GFP) (Helker et al., 2013), Tg(krt5:Lifeact-Ruby), and Tg(krt5:Myl12.1-EGFP) (van Loon et al., 2020). Tg(krt5:Gal4/+;UAS:Lifeact-GFP/+) zebrafish were incrossed or outcrossed to WT and screened for brightness on the day of imaging using a fluorescence dissecting microscope. For Airyscan microscopy, Tg(krt5:Myl12.1-EGFP) zebrafish were incrossed and injected with krt5:Lifeact-Ruby and krt5:Myl12.1-EGFP plasmids to improve brightness. All experimental procedures were approved by the Chancellor’s Animal Research Care Committee at the University of California, Los Angeles.

### Plasmids

Previously characterized plasmids in this paper include krt5:Myl12.1-EGFP (van Loon et al., 2020) and UAS:mRuby-PH-PLC (Jiang et al., 2019). krt5-Myh9a-mCherry was constructed using the Gateway-based Tol2-kit (Kwan et al., 2007). The following vectors used to construct krt5-Myh9a-mCherry have previously been described: p5E-krt5 (Rasmussen et al., 2015), p3E-mCherrypA (Kwan et al., 2007), and pDestTol2pA2 (Kwan et al., 2007). The Myh9a coding sequence was cloned from a cDNA library of 5dpf zebrafish larvae using the following primers: Forward: 5’-GGGGACAAGTTTGTACAAAAAAGCAGGCTATATGTCAGACGCAGAGAAGTTC-3’; Reverse: 5’-GGGGACCACTTTGTACAAGAAAGCTGGGTCTTACTCAGGAGTTGGCTCG-3’.

For transient transgene expression, ~5 nL plasmid (~25 ng/μL) was injected into single cell zebrafish embryos.

### Microscopy

Live fluorescent images and videos of periderm cells were acquired on a Zeiss LSM800 confocal microscope. Images were acquired with Zeiss Zen Blue software using an EC Plan-Neofluar 40×/1.30 oil DIC M27 objective with 2–3× digital zoom. Optimal resolution and Z-stack intervals were set using Zen software, except for videos for which a Z-stack interval of 0.75 μm was used to improve imaging speed. During imaging, zebrafish slide chambers were mounted on a heated stage set to 28°C. The x-y position and z-stack were occasionally adjusted during time-lapse imaging to keep the cells of interest in the frame. For longitudinal experiments between 48-96 hpf, zebrafish were rescued from mounting agarose each day after imaging using forceps, then placed in separate petri dishes for mounting and imaging on subsequent days.

Airyscan microscopy was performed on a Zeiss LSM 880 confocal microscope with Airyscan in the Broad Stem Cell Institute Research Center/Molecular, Cell and Developmental Biology microscopy core at UCLA. Images were acquired with Zeiss Zen Black software using an Plan-Apochromat 63x/1.4 Oil DIC M27 objective with 2–5× digital zoom. After acquisition, Airyscan processing was performed with the default settings on Zen Black.

To ablate periderm cells expressing Lifeact-GFP, we adapted a previously described method (O’Brien et al., 2009a; van Loon et al., 2020). Videos of cell stretching by periderm cell ablation were acquired using Zeiss Zen Black Software on a Zeiss LSM 880 multiphoton microscope using an EC Plan-Neofluar 40×/1.30 oil DIC M27 objective and a Coherent Chameleon Ultra II laser at a wavelength of 813 nm. A 488-nm laser was used to find and focus on the cell surface at 250× digital zoom, and the cell was then exposed to 813 nm laser illumination for 3–4 s at 5–6% laser power using “live” scanning.

### Drug Treatment

(-)-Blebbistatin (Cayman Chemical) was dissolved in DMSO (Fisher Scientific). Treatment solutions were made with Ringer’s Solution and included the inhibitor, or equivalent concentration of DMSO (≤1%), as well as up to 0.4 mg/mL MS-222 (Sigma). Zebrafish larvae were exposed to the treatment solution for the specified period of time, then mounted in agarose and immersed in the same solution. For treatments longer than 2 hours, larvae were initially exposed to a treatment solution without MS-222 and then transferred to a similar solution containing up to 0.4 mg/mL MS-222 ≥30min prior to imaging. For longitudinal experiments with blebbistatin, fish were first mounted in agarose and imaged, then rescued from agarose using forceps and exposed to treatment solutions. Approximately 30 minutes after exposure to treatment solutions, zebrafish were again mounted in agarose and slide chambers were filled with treatment solution. Zebrafish were imaged again after 1-hour exposure to treatment solutions.

### Image Analysis and Statistics

All statistical testing was performed using RStudio (RStudio, Inc.). Data distributions were assessed for normality using the Shapiro-Wilk test and visually inspected using Q-Q plots. The appropriate parametric or non-parametric tests were then selected based on the normality of the data distributions being compared.

Microridge analysis was performed using a custom Python script. Images of periderm cells were sum-projected and smoothened with a Gaussian filter. Pixel intensities were then normalized based on the modality of their intensity distribution. Unimodal distributions were normalized to the full width at the half maximum, while bimodal distributions were normalized to values between both maxima. Images were then processed with a Hessian filter, thresholded and skeletonized. Vectorized skeletons were smoothened and fitted to a normalized cell image to produce vectorized microridge lines. Distances between microridges and microridge orientations were then calculated. Microridge alignment domains were calculated by interpolating Q-tensor (**Q** = **v** ⊗ **v** - ½**I**, where **v** is a unit tangent vector and **I** is a unit tensor).

Image management for presentation was performed using FIJI (Schindelin et al., 2012). The brightness and contrast of images were adjusted for the purpose of presentation. All movies were stabilized for presentation and analysis purposes using the Image Stabilizer FIJI plugin (“Kang Li @ CMU - Image Stabilizer Plugin for ImageJ,” n.d.).

Microridge fusion and fission events were identified manually using the FIJI Multi-point tool. To measure distances from NMII contractions to fusion and fission events, NMII images were smoothened and contractions were automatically thresholded in FIJI with the Triangle method, then distances were measured between microridge rearrangement events and the edge of the nearest contraction using ‘rgeos’ and ‘sp’ R packages.

## SUPPLEMENTAL FIGURE LEGENDS

**Fig S1.**
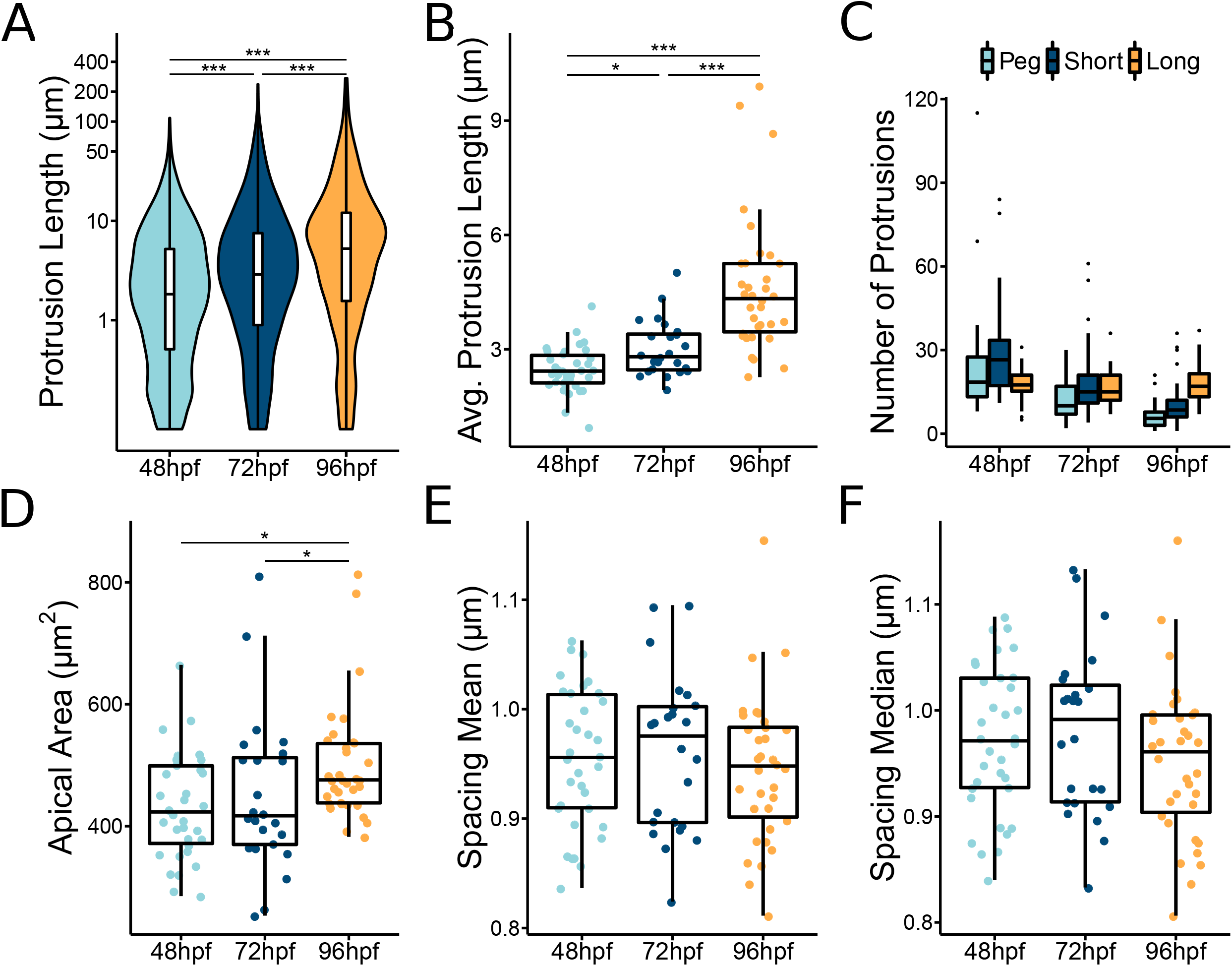
Additional quantification of morphological changes in maturing microridges. A) Violin and box-and-whisker plot of protrusion length for periderm cells at the specified stage. 48 hpf, n=34 cells from 12 fish; 72 hpf, n=24 cells from 10 fish; 96 hpf, n=34 cells from 15 fish. P<2.2×10^−16^, Kruskal-Wallis test followed by Dunn test with Benjamini-Hochberg p-value adjustment: 48-72 hpf, P=1.36×10^−14^; 48-96 hpf, P=2.67×10^−59^; 72-96 hpf, P=6.08×10^−14^ B) Dot and box-and-whisker plot of average protrusion length on periderm cells at the specified stage. 48 hpf, n=34 cells from 12 fish; 72 hpf, n=24 cells from 10 fish; 96 hpf, n=34 cells from 15 fish. P=1.81×10^−10^, Kruskal-Wallis test followed by Dunn test with Benjamini-Hochberg p-value adjustment: 48-72 hpf, P=0.019; 48-96 hpf, P=8.60×10^−11^; 72-96 hpf, P=3.05×10^−4^ C) Box-and-whisker plot of protrusion number distributed among pegs (<0.75μm), short microridges (0.75-5μm), and long microridges (>5μm) on periderm cells at the specified stage. 48 hpf, n=34 cells from 12 fish; 72 hpf, n=24 cells from 10 fish; 96 hpf, n=34 cells from 15 fish. Two-way ANOVA with interaction: hpf, P=3.12×10^−12^; protrusion type, P=4.84×10^−3^; hpf-protrusion type interaction, P=3.03×10^−5^. D) Dot and box-and-whisker plot of periderm cell apical area at the specified stage. 48 hpf, n=34 cells from 12 fish; 72 hpf, n=24 cells from 10 fish; 96 hpf, n=34 cells from 15 fish. P=0.011, Kruskal-Wallis test followed by Dunn test with Benjamini-Hochberg p-value adjustment: 48-72 hpf, P=0.722; 48-96 hpf, P=0.014; 72-96 hpf, P=0.041. E) Dot and box-and-whisker plot of microridge spacing mean for periderm cells at the specified stage. 48 hpf, n=34 cells from 12 fish; 72 hpf, n=24 cells from 10 fish; 96 hpf, n=34 cells from 15 fish. P=0.308, one-way ANOVA. F) Dot and box-and-whisker plot of microridge spacing median for periderm cells at the specified stage. 48 hpf, n=34 cells from 12 fish; 72 hpf, n=24 cells from 10 fish; 96 hpf, n=34 cells from 15 fish. P=0.569, one-way ANOVA. ‘*’ p ≥ 0.05 and ‘***’ p ≥ 0.001. For box-and-whisker plots, middle box line is the median, and lower and upper ends of boxes are 25th and 75th percentiles, respectively.

## VIDEO LEGENDS

**Video 1. Microridge fusion and fission diminish as microridge patterns mature** 10-minute time-lapse videos with 30-second intervals of periderm cells expressing Lifeact-GFP in zebrafish at the indicated developmental stage. Microridge fusion and fission attenuate as microridges become longer and more aligned at each stage. Orange circles show locations of microridge fusions. Blue circles show locations of microridge fissions. Scale bar: 10μm.

**Video 2. Microridges fuse and fission** 4.5-minute time-lapse videos with 30-second intervals of periderm cells expressing Lifeact-GFP in 48hpf zebrafish. White arrowheads show locations of microridge fusion and fission events. Scale bar: 1μm.

**Video 3. Microridge fusion and fission reflect fusion and fission of the plasma membrane** 10-minute time-lapse video with 30-second intervals of periderm cells expressing fluorescent reporters for actin (Lifeact-GFP) and membrane (mRuby-PH-PLC) on 48hpf zebrafish. Microridge fusion (yellow arrowhead) and fission (white arrowhead) in the actin channel are mimicked by fission and fusion of projections in the membrane channel. Time-lapse frames are sum projection images. Scale bar: 1μm.

**Video 4. Rapid cell shape changes do not induce microridge fusion and fission** 60-minute time-lapse video with 1-minute intervals of periderm cells expressing Lifeact-GFP on 72hpf zebrafish. Time-lapse begins immediately after laser ablation of periderm cells on either side of the cell of interest. The cell of interest rapidly elongates between the two wounds, but does not increase fusion and fission events. Orange circles show locations of microridge fusions. Blue circles show locations of microridge fissions. Microridge rearrangements occurred at a rate of 0.00393 events/μm · min over the course of the video. Scale bar: 10μm.

**Video 5. NMII contractions correlate with microridge fusion and fission** 10-minute time-lapse video with 30-second intervals of periderm cells expressing fluorescent reporters for actin (Lifeact-mRuby) and NMII (Myl12.1-EGFP) on 48hpf zebrafish. Microridges fuse near sites of intensifying myosin fluorescence signal (yellow arrowheads) and fission near sites of diminishing myosin fluorescence signal (white arrowhead). Scale bar: 1μm.

**Video 6. Short-term NMII inhibition reduces microridge fusion and fission** 10-minute time-lapse video with 30-second intervals of periderm cells expressing Lifeact-GFP on 49hpf zebrafish after 1-hour treatment with 1% DMSO or 50μM blebbistatin. Microridge fusion and fission decrease in periderm cells after 1-hour treatment with blebbistatin. Orange circles show locations of microridge fusions. Blue circles show locations of microridge fissions. Scale bar: 5μm.

**Video 7. NMII minifilaments coordinate peg dynamics and microridge fusion, fission, and spacing** 9-minute time-lapse videos with 1-minute intervals of periderm cells expressing fluorescent reporters for actin (Lifeact-mRuby) and NMII (Myl12.1-EGFP). NMII minifilaments appear as two green puncta. Different NMII-mediated events indicated by title cards. “Bridges” of one or two NMII minifilaments attach to pegs as they appear, and occasionally pull them toward one another. NMII minifilaments connect two microridge ends and fuse them into a longer microridge. NMII minifilaments oriented perpendicular to a microridge in the x-y plane sever a microridge. Finally, NMII minifilament “bridges” connecting two adjacent microridges contract to pull the microridges closer together, and allow microridges to drift further apart as they disappear. Scale bar: 1μm.

## ACKNOWLEDGMENTS

We thank Sally Horne-Badovinac, Yasuko Inaba, and Kadidia Pemba Adula for comments on the manuscript, Son Giang and Linda Dong for excellent fish care, and Nat Prunet in the BSCRC/MCDB microscopy core for help with Airyscan microscopy. This work was funded by National Institutes of Health grants R21EY024400 and R01GM122901 to A. Sagasti, and by the Biotechnology and Biological Sciences Research Council of UK grants BB/P01190X and BB/P006507 to A.B. Goryachev. A.P. van Loon was supported by the Ruth L. Kirschstein National Research Service Award (GM007185).

## Notes

### Competing Interest Statement

The authors have declared no competing interest.

### Summary of Updates

This version of the manuscript has been revised to update the organization of the text and make minor changes to figures and videos.

